# riboviz 2: A flexible and robust ribosome profiling data analysis and visualization workflow

**DOI:** 10.1101/2021.05.14.443910

**Authors:** Alexander L. Cope, Felicity Anderson, John Favate, Michael Jackson, Amanda Mok, Anna Kurowska, Emma MacKenzie, Vikram Shivakumar, Peter Tilton, Sophie M. Winterbourne, Siyin Xue, Kostas Kavoussanakis, Liana F. Lareau, Premal Shah, Edward W.J. Wallace

## Abstract

**Motivation:** Ribosome profiling, or Ribo-seq, is the state of the art method for quantifying protein synthesis in living cells. Computational analysis of Ribo-seq data remains challenging due to the complexity of the procedure, as well as variations introduced for specific organisms or specialized analyses. Many bioinformatic pipelines have been developed, but these pipelines have key limitations in terms of functionality or usability.

**Results:** We present riboviz 2, an updated riboviz package, for the comprehensive transcript-centric analysis and visualization of Ribo-seq data. riboviz 2 includes an analysis workflow built on the Nextflow workflow management system, combining freely available software with custom code. The package is extensively documented and provides example configuration files for organisms spanning the domains of life. riboviz 2 is distinguished by clear separation of concerns between annotation and analysis: prior to a run, the user chooses a transcriptome in FASTA format, paired with annotation for the CDS locations in GFF3 format. The user is empowered to choose the relevant transcriptome for their biological question, or to run alternative analyses that address distinct questions. riboviz 2 has been extensively tested on various library preparation strategies, including multiplexed samples. riboviz 2 is flexible and uses open, documented file formats, allowing users to integrate new analyses with the pipeline.

**Availability:** riboviz 2 is freely available at github.com/riboviz/riboviz.

**Supplementary information:** 

## 1 Introduction

Ribo-seq allows researchers to quantify the “translatome” of actively translated RNAs in cells (Ingolia *et al*., 2009). Ribo-seq combines high-throughput sequencing with nuclease footprinting of ribosomes to identify the location of ribosomes across the transcriptome at codon-level resolution. Ribo-seq is often combined with RNA-seq to quantify post-transcriptional regulation and also provides quantitative information on the movement of ribosomes along an mRNA, allowing mechanistic insight into translation. The emergence of Ribo-seq has necessitated the development of pipelines to enable data pre-processing, read mapping, gene-specific quantification, and other downstream analyses (Li *et al*., 2020). Ribo-seq data presents unique challenges relative to other RNA sequencing methods, and analysis remains specialized. To address these challenges, we previously developed riboviz, an analysis and visualization framework that provided users with the ability to analyze their own private data, as well as a web-based tool for the comparison of Ribo-seq measurements (Carja *et al*., 2017). Here, we present a significantly expanded and reworked version of riboviz: riboviz 2 (Wallace *et al*., 2020).

## 2 Methods

The riboviz 2 pipeline is implemented via Nextflow (Di Tommaso *et al*., 2017; Jackson *et al*., 2021), with run-specific parameters specified by the user in a YAML-format configuration file. The configuration file specifies the input FASTQ and annotation files, as well as all run-specific parameters (see Documentation at github.com/riboviz/riboviz). riboviz 2 invokes both publicly available tools (e.g. cutadapt (Martin, 2011), HISAT2 (Kim *et al*., 2015), UMI-Tools (Smith *et al*., 2017)), and custom Python and R scripts for data parsing and visualization. Running the workflow via Nextflow and using a single configuration file avoids the need to configure and run each tool individually, facilitates reproducible and transparent analyses, and allows the pipeline to run on various computing systems. riboviz 2 is capable of processing multiple measurements from an experiment, including multiplexed libraries, in a single command-line call.

The riboviz 2 workflow (Fig. 1A) starts with pre-processing of Ribo-seq data, including adapter trimming and removing reads mapping to user-supplied contaminant sequences such as rRNA. Following pre-processing, the remaining reads are aligned to the relevant sequences as defined by the provided FASTA and GFF3 files. Due to differences in ribosome structure between Archaea, Bacteria, and Eukarya, the appropriate strategy for assigning Ribo-seq reads to the codon at the ribosomal active site varies between species. Reads from eukaryotes are usually assigned to an A-site codon at a fixed displacement from their 5’ end, but assigning relative to the 3’ end is more accurate in bacteria (Woolstenhulme *et al*., 2015; Mohammad *et al*., 2019). riboviz 2 allows the user to map relative to either end of the read by specifying the displacement separately for each desired read length.

**Fig. 1.**
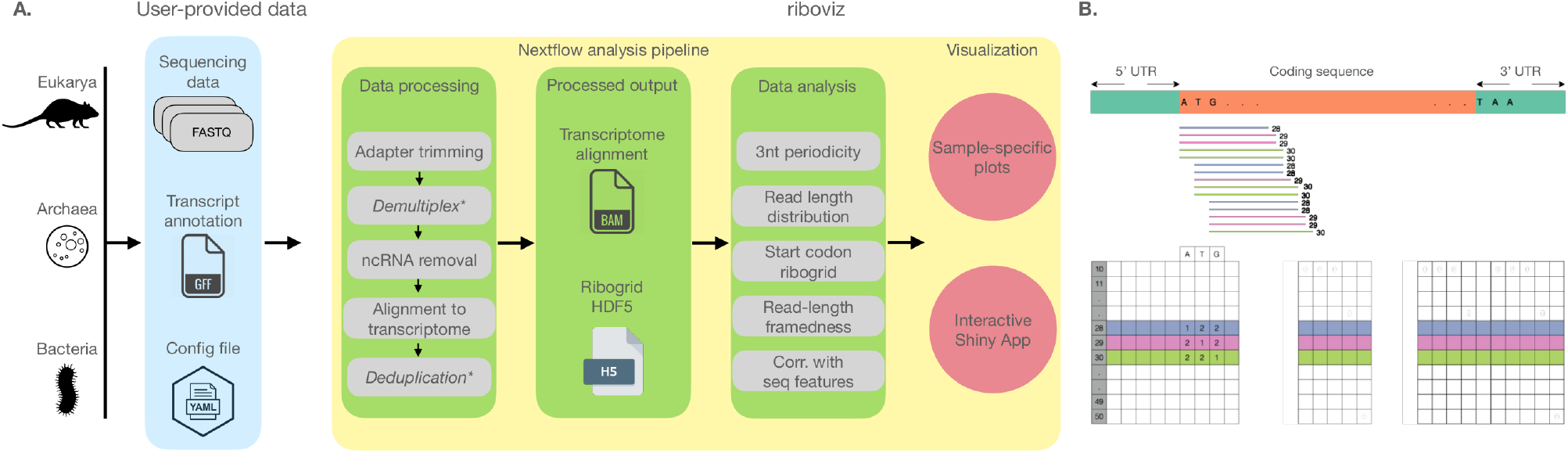
riboviz pipeline and data structures. A. riboviz takes in user-provided sequencing, transcript annotation, and configuration files, processes the datasets, and generates two major outpust - transcriptome-specific BAM file and a ribogrid file. These outputs are used to generate sample-specific analyses and summaries, which can be visualized as both static figures and in an interactive R/Shiny application. B. Structure of the ribogrid file format. Ribogrid is a complete representation of transcript-specific ribosome-footprint data in an H5 file-format. Each row indicates reads of a particular length and each column indicates the position of the 5’-end of a footprint. *optional.

riboviz 2 provides outputs typical to Ribo-seq in standard file formats, including the aligned reads in BAM format, and text files with data such as three-nucleotide periodicity and read counts by read length. We also provide an intermediate data file in H5 format that contains read counts organized into matrices by both 5’ position and read length. These counts are a sufficient statistic for most downstream analyses, in that the only information used from the raw alignments is the count by both position and length. The H5 file contains one “aligned read count matrix” per transcript per sample. The riboviz 2 H5 file format is documented and accessor functions for its contents are provided. These enable the future addition of custom analysis functions. The H5 format balances user accessibility with a compact file size.

True to its name, riboviz 2 automatically outputs visualizations that include plots of three-nucleotide periodicity and read-length distributions. These visualizations are commonly used in publications describing Ribo-seq experiments, both for quality control to confirm that the experiment successfully recovered ribosome footprints, and as a valuable tool for analysis. riboviz 2 directly visualizes the “aligned read count matrix” that is stored in the H5 file, by showing a heatmap of the footprint counts arranged by both 5’ position and read length (Fig. 1B). These “Ribogrid’ ’ plots are a rich way to read out mechanistic details of Ribo-seq data such as read frame (Lareau *et al*., 2014).

## 3 New Features and Advantages

### Flexibility across organisms

riboviz 2 can be used on any organism across the three major domains of life (Archaea, Bacteria, and Eukarya) for which a transcriptome FASTA and GFF3 file can be constructed. The user is responsible for supplying a FASTA file appropriate to their biological question, for example using a published annotation to define spliced transcripts including UTRs, or a “padded ORFeome” that contains fixed-width extensions to a set of ORFs of interest. The user must also supply a file in GFF3 format that specifies the positions of the ORFs of interest within the transcripts. This flexibility distinguishes riboviz 2 from Ribo-seq bioinformatics pipelines that are organism or database specific, limiting their use to the scientific community (Wang *et al*., 2018; Perkins *et al*., 2019). We maintain a repository of example analyses, including transcriptome and contamination files, that span the major domains of life, at github.com/riboviz/example-datasets.

### Usability

All the parameters for all steps of a riboviz 2 analysis are specified in a single YAML-format configuration file, with thorough documentation. We include configuration files for diverse datasets in our repository github.com/riboviz/example-datasets. These files may be used to reproduce analyses, or adapted to analyzing new Ribo-seq datasets. Thus, no knowledge of Python or R is required to take advantage of the riboviz 2 functionality, unlike many other tools (Backman and Girke, 2016; Lauria *et al*., 2018). The full documentation of riboviz 2 can be found at github.com/riboviz/riboviz.

### Flexible end-to-end data processing workflow

riboviz 2 analyses Ribo-seq data from raw reads through to publication-quality figures. Few Ribo-seq analysis tools provide comprehensive data pre-processing (e.g. adapter trimming) or read alignment (Li *et al*., 2020). Following read alignment, riboviz 2 uniquely invokes an (optional) script to trim non-templated mismatches added by some viral reverse transcriptases (Wulf *et al*., 2019), which otherwise leads to inaccurate quantification of read frame. riboviz 2 is also more flexible in the varieties of library preparation methods that it can process. To the best of our knowledge, riboviz 2 is the only Ribo-seq pipeline which is pre-built to handle multiplexed libraries or unique molecular identifiers via UMI-Tools (Smith *et al*., 2017).

### Flexible and documented data outputs

A major goal of a Ribo-seq analysis pipeline is to enable further downstream analyses of Ribo-seq data, such as differential expression analysis and identification of ribosome pausing sites. riboviz 2 consolidates the data into outputs that are suitable for downstream analysis, such as the read count matrices H5 file. It aggregates raw counts per transcript into a format which can be used as input to tools such as DESeq2 (Love *et al*., 2014) and provides per-ORF translation TPM values.

Overall, riboviz 2 is a flexible and carefully engineered open-source workflow for Ribo-seq analysis and visualization that we hope will find broad use.

## Acknowledgements Funding

This work was supported by the Biotechnology and Biological Sciences Research Council [BB/S018506/1 to E.W.J.W.]; the Wellcome Trust [208779/Z/17/Z to E.W.J.W.]; the National Science Foundation [DBI 1936046 to P.S., DBI 1936069 to L.F.L.]; the National Institutes of Health [R35 GM124976 and subcontracts from R01 DK056645, R01 DK109714, R01 DK124369 to P.S.]; and start-up funds from the Human Genetics Institute of New Jersey at Rutgers University to P.S.

